# Spiking network optimized for noise robust word recognition approaches human-level performance and predicts auditory system hierarchy

**DOI:** 10.1101/243915

**Authors:** Fatemeh Khatami, Monty A. Escabí

## Abstract

The auditory neural code is resilient to acoustic variability and capable of recognizing sounds amongst competing sound sources, yet, the transformations enabling noise robust abilities are largely unknown. We report that a hierarchical spiking neural network (HSNN) trained to maximize word recognition accuracy in noise and multiple talkers approaches human-level performance. Intriguingly, comparisons with data from auditory nerve, midbrain, thalamus and cortex reveals that the organization and nonlinear transformations of the optimal network predict several properties of the ascending auditory pathway including a sequential loss of temporal resolution, increasing sparseness and selectivity. The optimal organizational scheme is critical for noise robustness since an identical network arranged to enable high information transfer does not predict auditory pathway organization and has substantially poorer performance. Furthermore, conventional linear and nonlinear receptive field-based models fail to achieve similar noise robust performance. The findings suggest that the auditory pathway hierarchy and its sequential nonlinear feature extraction computations may form a near optimal code capable of efficiently detecting sounds in noise impoverished conditions.

**Significance Statement:** The brain’s ability to recognize sounds in the presence of competing sounds or background noise is essential for everyday hearing tasks. How the brain accomplishes noise resiliency, however, is poorly understood. Using neural recording from the ascending auditory pathway and an auditory spiking network model trained for optimal sound recognition in noise we explore the computational strategies that enable noise robustness. Our results suggest that the hierarchical organization of the auditory pathway and the resulting nonlinear transformations may form a near optimal strategy that is essential for sound recognition in the presence of noise.

## Introduction

Being able to identify sounds in the presence of background noise is essential for everyday audition and vital for survival. Although several cortical mechanisms have been proposed to facilitate robust coding of sounds ^1,2^ it is presently unclear how the sequential organization of the ascending auditory pathway and the resulting nonlinear transformations contribute to robust sound recognition.

Several hierarchical changes in spectral and temporal selectivity are consistently observed in the ascending auditory pathway of mammals. Temporal selectivity and resolution change dramatically over more than an order of magnitude, from a high-resolution representation in the cochlea, where auditory nerve fibers synchronize to temporal features of up to ~1000 Hz, to progressively slower (limited to ~25 Hz) and coarser resolution representation as observed in auditory cortex ^3^. Furthermore, although changes in spectral selectivity can be described across different stages of the auditory pathway, and spectral resolution is somewhat coarser in central levels, changes in frequency resolution are somewhat more homogeneous and less dramatic ^4-6^. It is plausible that such hierarchical transforms across auditory nuclei are essential for feature extraction and ultimately high-level auditory tasks such as acoustic object recognition. Yet, it is unclear whether these sequential transformations comprise an optimal computational strategy for noise robust sound encoding. Here we report that the hierarchical organization of the auditory pathway and its sequential nonlinear feature extraction transformations form a near-optimal computation strategy for noise robust sound coding.

## Results

### Task optimized hierarchical spiking neural network predicts auditory system organization

We developed a physiologically motivated hierarchical spiking neural network (HSNN) and trained it on a behaviorally relevant word recognition task in the presence of background noise and multiple talkers. Like the auditory pathway, the HSNN receives frequency-organized input from a cochlear stage (Fig. 1**a**) and maintains its topographic (tonotopic) organization through a network of frequency organized integrate-and-fire spiking neurons (Fig. 1**b**). For each sound, such as the word “zero”, the network produces a dynamic spatio-temporal pattern of spiking activity (Fig. 1**b**, right) as observed for peripheral and central auditory structures ^7-9^. Each neuron is highly interconnected containing frequency specific and co-tuned excitatory and inhibitory connections ^10-13^ that project across six network layers (Fig. 1**b**). Converging spikes from neurons in a given layer (Fig 1**d**) are weighted by frequency localized excitatory and inhibitory connectivity functions and the resulting excitatory and inhibitory post-synaptic potentials are integrated by the recipient neuron (Fig. 1**d** and **e**, note the variable spike amplitudes). Output spike trains from each neuron are then weighted by connectivity function, providing the excitatory and inhibitory inputs to the next layer (Fig. 1**e, f**). The overall multi-neuron spiking output of the network (Fig. 1**b**, right) is then treated as a response feature vector and fed to a Bayesian classifier in order to identify the original sound delivered (Fig. 1**c**; see Methods).

**Figure 1.**
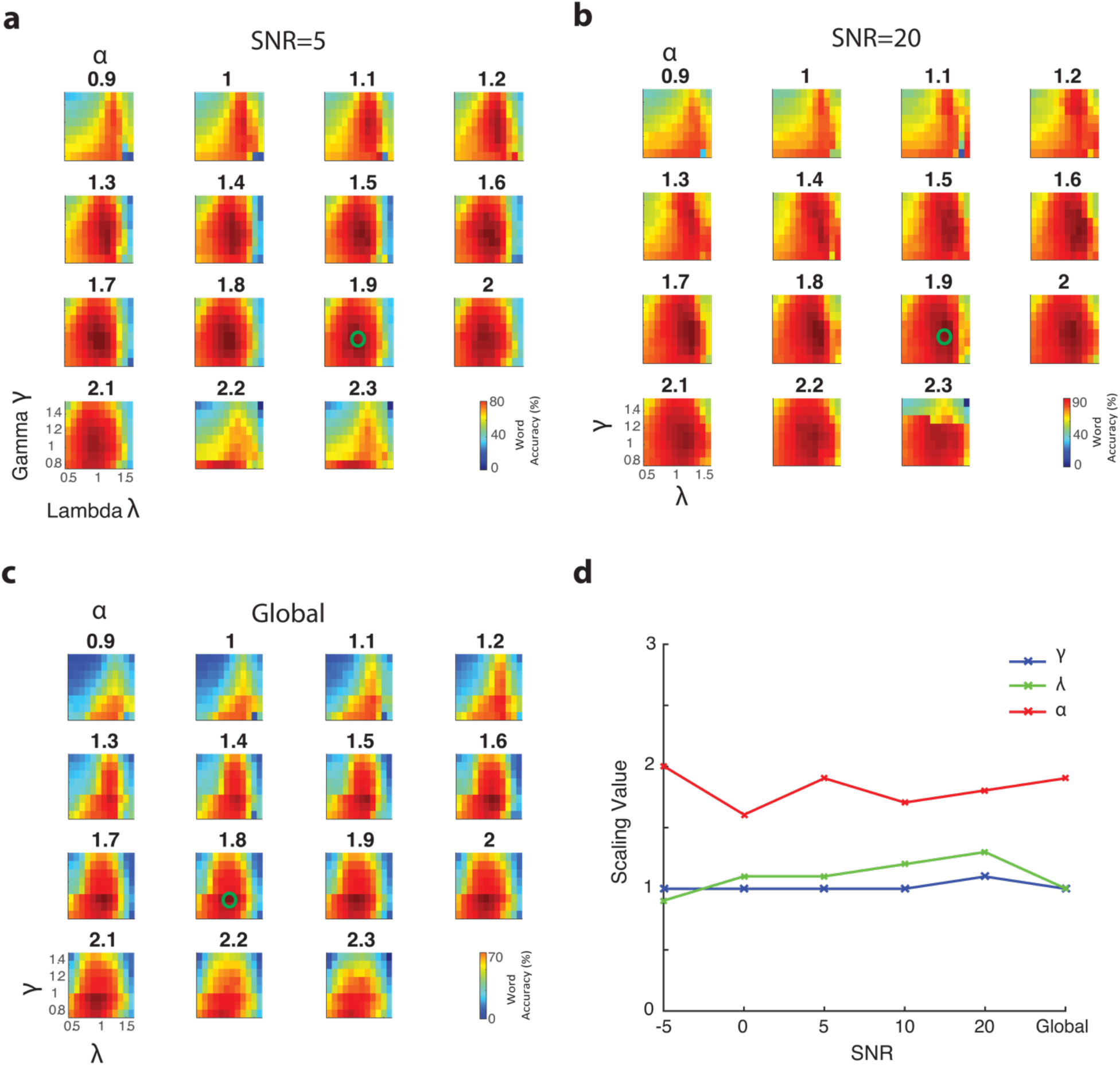
Auditory pathway *hierarchical spiking neural network* (HSNN) model. The model consists of a **(a)** cochlear model stage that transforms the sound waveform into a spectrogram (time vs. frequency), **(b)** a central hierarchical spiking neural network containing frequency organized spiking neurons and a **(c)** Bayesian classifier that is used to read the spatio-temporal spike train outputs of the HSNN. Each dot in the output represents a single spike at a particular time-frequency bin. **(d-f)** Zoomed in view of the HSNN illustrates the pattern of convergent and divergent connections between network layers for a single leaky integrate-and-fire (LIF) neuron. **(d-e)** Input spike trains from the preceding network layer are integrated with excitatory (red) and inhibitory (blue) connectivity weights that are spatially localized and model by Gaussian functions **(f)**. The divergence and convergence between consecutive layers is controlled by the connectivity width (SD of the Gaussian model, *σ_l_*). Each incoming spike generates excitatory and inhibitory post-synaptic potentials (EPSP and IPSP, red and blue kernels in **e**). The integration time constant (*τ_l_*) of the EPSP and IPSP kernels can be adjusted to control the temporal integration between consecutive network layers while the spike threshold level (*N_l_*) is independently adjusted to control the output firing rates and the overall neuron layer sensitivity. (**g, h**) Example cochlear model outputs and the corresponding multi-neuron spike train outputs of the HSNN under the influence of speech babble noise (at 20 dB SNR). **(g)** HSNN response pattern for one sample of the words *zero, six*, and *eight* illustrate output pattern variability that can be used to differentiate words. **(h)** Example response variability for the word *zero* from multiple talkers in the presence of speech babble noise (20 dB SNR).

Given that key elements of speech such as formants and phonemes have unique spectral and temporal composition that are critical for word identification ^14,15^, we first test how the spectro-temporal resolution and sensitivity of each network layer contribute to word recognition performance in background noise. We optimize the HSNN to maximize word recognition accuracy in the presence of noise and to identify the network organization of three key parameters that separately control the temporal and spectral resolution and the overall sensitivity of each network layer (*l*=1 … 6). The neuron time-constant (*τ_l_*), controls the temporal dynamics of each neuron element in layer *l* and the resulting temporal resolution of the output spiking patterns. The connectivity width (*σ_l_*) controls the convergence and divergence of synaptic connections between consecutive layers and therefore affects the spectral resolution of each layer. Since synaptic connections in the auditory system are frequency specific and localized ^13,16,17^ connectivity profiles between consecutive layers are modeled by a Gaussian profile of unknown connectivity width parameter ^18^ (Fig. 1**e**; specified by the SD, *σ_l_*). Finally, the sensitivity and firing rates of each layer are controlled by adjusting the spike threshold level (*N_l_*) of each IF neuron ^19^. This parameter controls the firing pattern from a high firing rate dense code as proposed for the auditory periphery to a sparse code as has been proposed for auditory cortex ^2,20^. Because temporal and spectral selectivities vary systematically and gradually across auditory nuclei^3,6,21^, we required that the network parameters vary hierarchically and smoothly from layer-to-layer according to (see Methods: Network Constraints and Optimization)

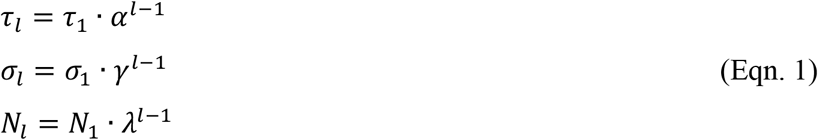

where *τ_l_*, *σ*_1_, and *N*_1_ are the parameters of the first network layer and are chosen so that first layer responses mimic activity in auditory nerve fibers (see Methods). The scaling parameters *α, λ*, and *γ* determine the direction and magnitude of layer-to-layer changes for each of the three neuron parameters. Scaling values greater than one indicate that the neuron parameter increases systematically across layers, a value of one indicates that the parameter is constant, while a value less than one indicates that the parameter value decreases systematically across layers.

The optimal network outputs preserve important time-frequency information in speech despite variability in the input sound. Sounds in the optimization and validation corpus consist of spoken words for digits from zero to nine from eight talkers (TI46 LDC Corpus ^22^, see Methods). As a task we require that the network identify the word (i.e., the digit) that is delivered as input (10 alternative forced choice task). Example cochlear model spectrograms and the network spiking outputs are shown in Fig. 1**g** and **h** for the words *zero, six*, and *eight* in the presence of speech babble noise (optimal outputs at SNR=20 dB). Analogous to auditory cortex responses for speech^7^, the network produces a distinguishable spiking output for each sound that reflects its spectro-temporal composition (Fig. 1**g**). Furthermore, when a single word is generated by different talkers in noise (SNR=20 dB) the network produces a relatively consistent firing pattern (Fig. 1**g**) such that the response timing and active neuron channels remain relatively consistent. For instance, a lack of activity is observed for neurons between ~2-4 kHz within the first ~100-200 ms of the sound for the word *zero* and several time-varying response peaks indicative of the vowel formants are observed for all three talkers (Fig. 1**h**).

To determine the network architecture required for optimal word recognition in noise and to identify whether such a configuration is essential for noise robust performance, we searched for the network scaling parameters (*α, λ*, and *γ*) that maximize the network’s word recognition accuracy in a ten-alternative forced choice task for multiple talkers (8) and in the presence of speech babble noise (signal-to-noise ratios, SNR=-5, 0, 5, 10, 15, 20 dB; see Methods). For each input sound, the network spike train outputs are treated as response feature vectors and a Bayesian classifier (Fig. 1**c**; see Methods) is used to read the network outputs and report the identified digit (*zero* to *nine*). The network word recognition accuracy is shown in Fig. 2 as a function of each of the network parameters (*α, λ*, and *γ*) and SNR (**a**, SNR=5 dB; **b**, SNR=20 dB; **c**, average accuracy across all SNRs). At each SNR the word recognition accuracy profiles are tuned with the scaling parameter (i.e., concave function) which enables us to find an optimal scaling parameters that maximizes the classifier performance. Regardless of the SNR the optimal HSNN parameters are relatively constant (Fig. 2**d**; tested between -5 to 20 dB) implying that the network organization is relatively stable and invariant of the SNR (Fig. 2**a**-**c**; **a**=5 dB SNR, **b**=20 dB SNR, **c**=average across all SNRs). Intriguingly, several functional characteristics of the optimal network mirror those observed in the auditory pathway. Like the ascending auditory pathway where synaptic potential time-constants vary from sub-millisecond in the auditory nerve to tens of milliseconds in cortex^13,23-25^, time constants scale in the optimal HSNN (global optimal *α =* 1.9) over more than an order of magnitude between the first and last layer (1.9^5^ = 24.8 fold increase between the first and last layer; ~0.5 to 12.5 ms) indicating that temporal resolution becomes progressively coarser in the deep network layers. By comparison, the optimal connectivity widths do not change across layers (*γ* = 1.0). This result suggests that for the optimal HSNN temporal resolution changes dramatically while spectral resolution remains relatively constant across network layers, mirroring changes in spectral and temporal selectivity observed along the ascending auditory pathway ^3-6^.

**Figure 2.**
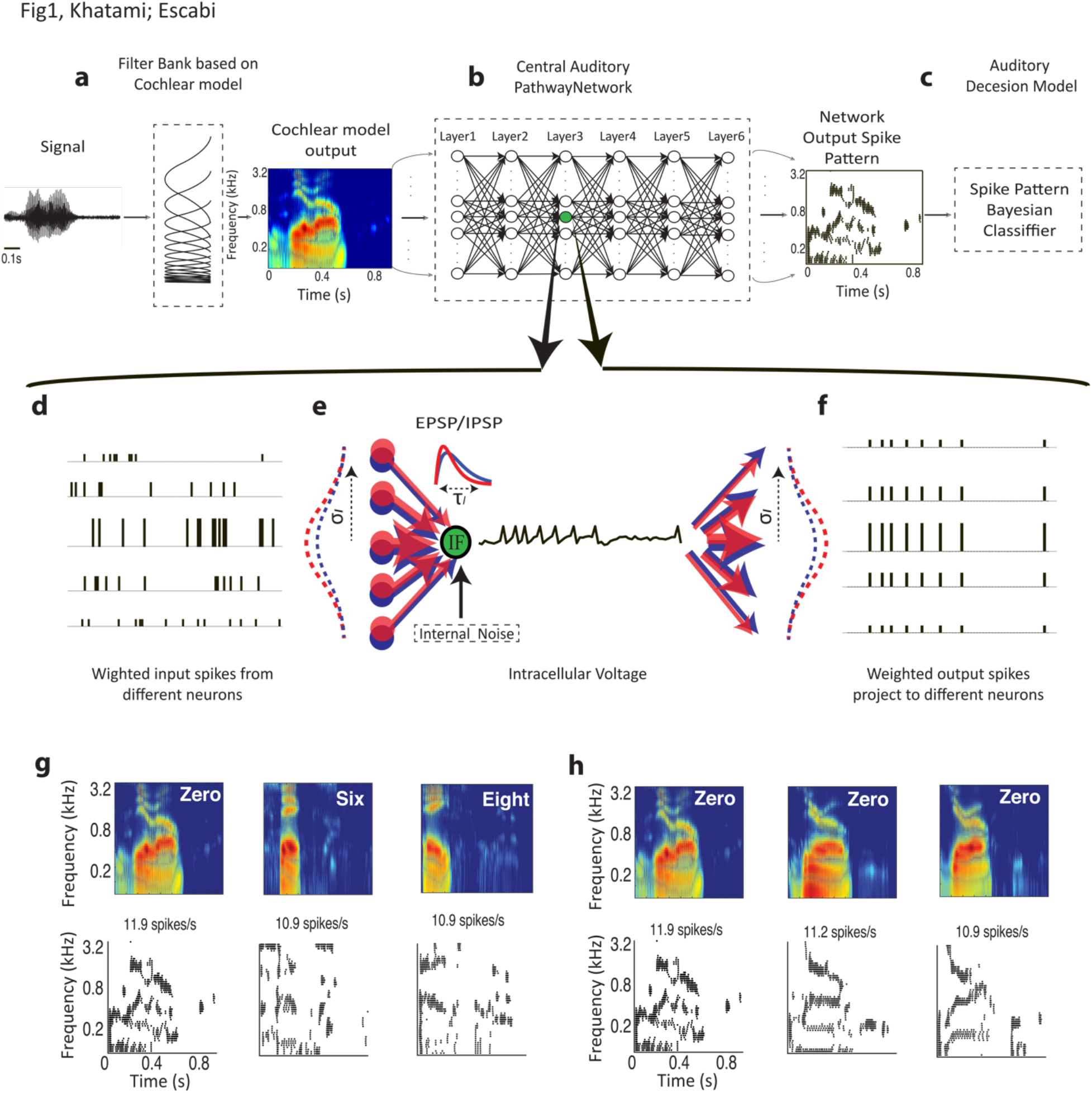
Hierarchical scaling is predicted by a global optimal solution that maximizes word recognition accuracy in the presence of background noise (-5, 0, 5, 10, 15 and 20 dB SNR). Crossvalidated word recognition accuracy (see Methods) is measured using the network outputs as a function of the three scaling parameters (*α, λ*, and *γ*). Word recognition accuracy curves are shown at 5 and 20 dB SNR (**a** and **b**, respectively) as well as for the global solution (**c**, average accuracy between -5 and 20 dB SNR). In all cases shown, word recognition accuracy curves are tuned for the different scaling parameters and exhibit a similar optimal solution (green circles). **(d)** The optimal scaling parameters are relatively stable across SNRs and similar to the solution that maximize average performance across all SNRs (optimal solution *α =* 1.9*, λ =* 1.0, and γ=1.0).

The scaling parameters of the optimal HSNN indicate a substantial loss of temporal (*α =* 1.9) and no change in connectivity resolution (*γ* = 1.0) across network layers. This prompted us to ask how feature selectivity changes across the network layers and whether a sequential transformation in spectral and temporal selectivity is essential for optimal word recognition in noise. To quantify the sequential transformations in acoustic processing, we first measure the spectro-temporal receptive fields (STRFs) of each neuron in the network (see Methods). Example STRFs are shown for two selected frequencies across the six network layers (Fig. 3 **a**; best frequency = 1.5 and 3 kHz). As a comparison, example STRFs from the auditory nerve (AN) ^26^, midbrain (inferior colliculus, IC) ^5^, thalamus (MGB) and primary auditory cortex (A1) ^6^ of cats are shown in Fig. 3**e**. Like auditory pathway neurons, STRFs from the optimal HSNN contain excitatory domains (red) with temporally lagged and surround inhibition/suppression (blue) along the frequency dimension (Fig. 3**a**). Furthermore, STRFs are substantially faster in early network layers lasting only a few milliseconds and mirroring STRFs from the auditory nerve, which have relatively short latencies and integration times. STRFs have progressively longer integration times (paired t-test with Bonferroni correction, p<0.01; Fig. 3**b**) and latencies (paired t-test with Bonferroni correction, p<0.01; Fig. 3**c**) across network layers, while bandwidths increase only slightly from the first to last layer (paired t-test with Bonferroni correction, p<0.01; Fig. 3**d**). These sequential transformations mirror changes in temporal and spectral selectivity seen between the auditory nerve, midbrain, thalamus and ultimately auditory cortex (Fig. 3**e-h**). As for the auditory network model, integration times (Fig. 3**f**) and latencies (Fig. 3**g**) increase systematically and smoothly (paired t-test with Bonferroni correction, p<0.01) while bandwidths show a small but significant increase between the auditory nerve and cortex (paired t-test with Bonferroni correction, p<0.01), analogous to results from the computational network. Although the network trends mirror changes in spectral and temporal selectivity seen between auditory nerve and cortex, auditory receptive fields tend to be somewhat slower and narrower than the network. Such disparities may partly be attributed to mechanisms not included in the HSNN such as descending feedback ^27^, synaptic and dendritic nonlinearities ^28^ and adaptive mechanisms such as spike time dependent plasticity, synaptic depression, and gain normalization ^1,29^.

**Figure 3.**
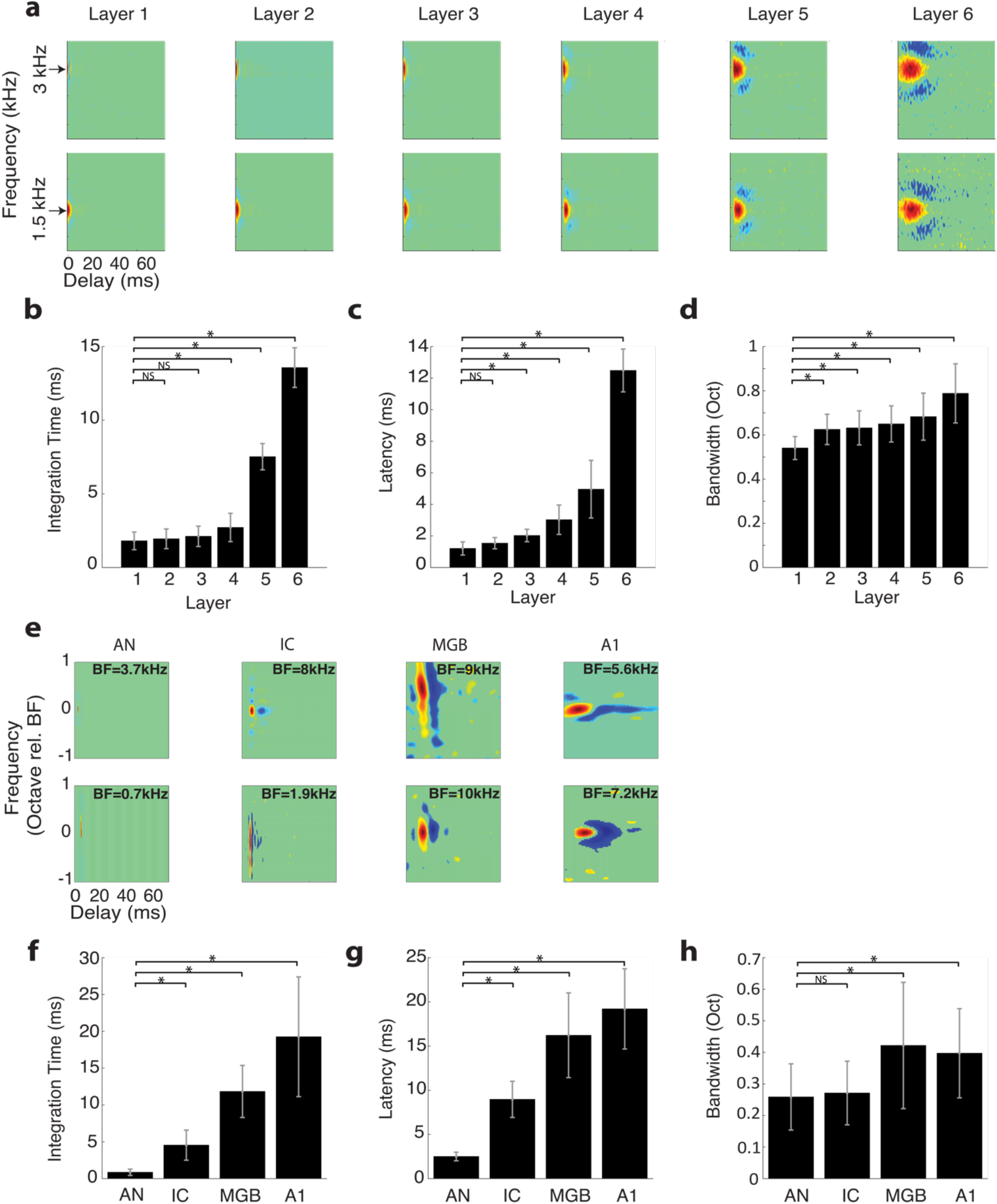
Receptive field transformations of the optimal HSNN predicts transformations observed along the ascending auditory pathway. **(a)** Example spectro-temporal receptive field (STRF) measured for the optimal network change systematically between consecutive network layers. All STRFs are normalized to the same color scale (red=increase in activity or excitation; blue=decrease in activity or inhibition/suppression; green tones=lack of activity). In the early network layers STRFs are relatively fast with short duration and latencies, and relatively narrowly tuned. STRFs become progressively slower, slightly broader, and have longer and more varied patterns of inhibition across the network layers, mirroring changes in spectral and temporal selectivity observed in the ascending auditory pathway. The measured **(b)** integration times, **(c)** latencies, and **(d)** bandwidths increase across the six network layers. **(e)** Examples STRFs from the auditory nerve (AN)^26^, inferior colliculus (IC)^5^, thalamus (MGB) and primary auditory cortex (A1)^6^ become progressively longer and have progressively more complex spectro-temporal sensitivity along the ascending auditory pathway. Average integration times **(f)**, latencies **(g)** and bandwidths **(h)** between AN and A1 follow similar trends as the optimal HSNN **(b-d)**. Asterisks (*) designate significant comparisons (t-test with Bonferroni correction, p<0.01) relative to layer 1 for the optimal network **(b-d)** or relative to the auditory nerve for the neural data **(f-h)** while error bars designate SD.

### Hierarchical and nonlinear transformations enhance robustness

It is intriguing that the hierarchical loss of temporal and spectral resolution in the optimal network mirror changes in selectivity observed in the ascending auditory system, as this ought to limit the transfer of acoustic information across the network. One plausible hypothesis is that such a sequential decrease in resolution is necessary to extract invariant acoustic features in speech while rejecting noise and fine details in the acoustic signal that may contribute in a variety of hearing tasks (e.g., spatial hearing, pitch perception etc.), but ultimately don’t contribute to speech recognition performance. This may be expected since human listeners require a limited set of temporal and spectral cues for speech recognition ^14,15^ and can achieve high recognition performance even when spectral and temporal resolution is degraded ^30,31^. We thus tested the above hypothesis by comparing the optimal network performance against a high-resolution network that lacks scaling (*α =* 1, *λ* = 1, and *γ* = 1) and for which we expect a minimal loss of acoustic information across layers. Unlike the optimal network, STRFs from the high-resolution network are relative consistent and change minimally across layers (Supplemental Data, Fig. 1**S**), which supports the idea that spectrotemporal information propagates across the high-resolution network with minimal processing.

Figure 4 illustrates how the optimal HSNN accentuates critical spectral and temporal cues necessary for speech recognition while the high-resolution network fails to do the same. Example Bayesian likelihood time-frequency histograms (average firing probability across all excerpts of each sound at each time-frequency bin) measured at 5 dB SNR are shown for the words “three”, “four”, “five” and “nine” for both the high-resolution (Fig. 4**a**) and optimal (Fig. 4**b**) HSNN along with selected spiking outputs from a single talker. Intriguingly, the Bayesian likelihood for the high-resolution network are highly blurred in both the temporal and spectral dimensions and have similar structure for the example words (Fig. 4**a**, right panels). This is also seen in the individual network outputs where the high-resolution network produces a dense and saturated firing pattern (Fig. 4**a**) that lacks the detailed spatio-temporal pattern seen in the optimal HSNN (Fig. 4**b**). The optimal HSNN preserves and even accentuates key acoustic elements such as temporal transitions for voice onset timing and spectral resonances (formants) while simultaneously rejecting and filtering out the background noise (Fig. 4**b**, right panels).

**Figure 4.**
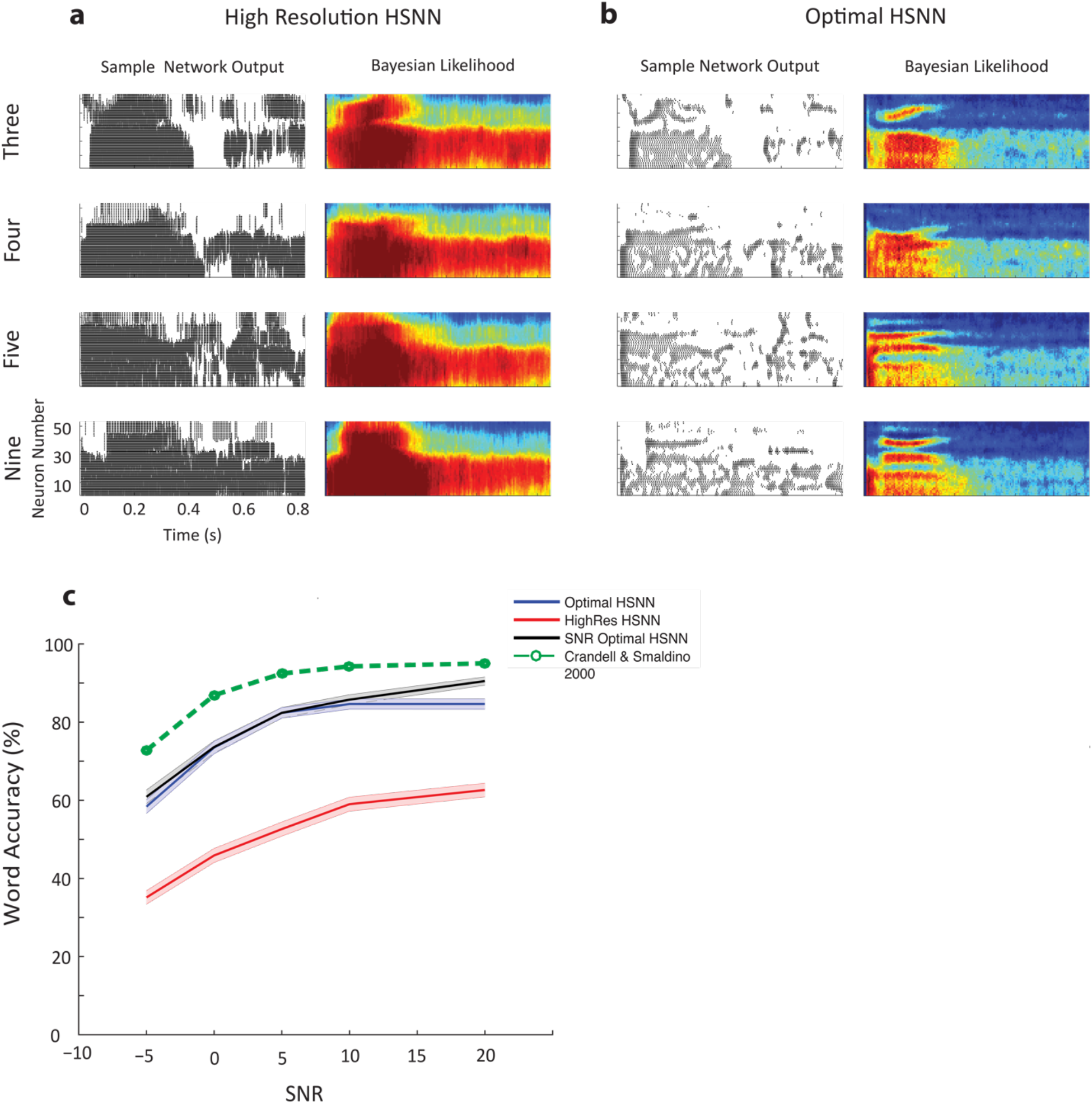
Optimal HSNN outperforms a high-resolution HSNN designed to preserve incoming acoustic information. Sample network spike train outputs and Bayesian likelihood histograms for the words *three*, *four*, *five*, and *nine* are shown for the **(a)** high-resolution and **(b)** optimal HSNN at 5 dB SNR. The Bayesian likelihood histograms correspond to the average probability of firing at each time-frequency bin for each digit (averaged across all trials and talkers). The firing patterns and Bayesian likelihood of the high-resolution network are spatio-temporally blurred compared to the hierarchical network. **(b)** Details such as spectral resonances (e.g., formants) and temporal transitions resulting from voicing onset are accentuated in the hierarchical network output. **(c)** The optimal HSNN (maximize performance across all SNRs) outperforms the high-resolution network in the word recognition task at all SNRs tested (blue=optimal; red=high-resolution) with an average accuracy improvement of 25.6 %. The optimal HSNN word recognition accuracy also closely matches the performance when the network is optimized and tested individually at each SNR (black, SNR optimal HSNN) indicative of a stable network representation. Finally, the optimal HSNN is within ~10% of human performance in a similar word recognition task (dotted-green curve ^32^).

We next compared the performance of the HSNN models to human subjects in an isolated monosyllabic word recognition task in speech babble noise ^32^. The word recognition accuracy of the optimal HSNN approaches human performance and is significantly higher than the high-resolution network for all of the SNRs tested (Fig. 4 **c**; green=human subjects^32^; p<0.001, t-test with Bonferroni correction). On average there is a 27.6 % improvement in the word accuracy rates for the optimal HSNN over the high-resolution HSNN. We also compared the accuracy of the optimal HSNN with the accuracy of a HSNN that was optimized individually at each SNR (SNR-optimal HSNN). The accuracy of the SNR-optimal HSNN was not significantly different from the optimal HSNN (p<0.05, t-test) which suggest that the optimal solution produces a stable noise robust representation. Furthermore, the optimal HSNN is on average within 11.5% of human performance in an isolated word recognition task and follows a similar performance trend across signal-to-noise ratios (Fig. 4**c**) ^32^.

To characterize the neural transformations enabling noise robust coding, we examine how acoustic information propagates and is transformed across sequential network layers. For each layer, the spike train outputs are first fed to the Bayesian classifier in order to measure sequential changes in word recognition accuracy. In the optimal HSNN, word recognition accuracy systematically increases across layers with an average improvement of 15.5% between the first and last layer when tested at 5 dB SNR (p<0.001, t-test; Fig. 5**a**, blue; 13.7% average improvement across all SNRs). By comparison, for the high-resolution HSNN, performance degrades sequentially across layers with an average decrease of 19.8% between the first and last layer (p<0.001, t-test; Fig. 5**a**, red; 18.1 % average reduction across all SNRs). Thus, the optimal HSNN is capable of sequentially extracting high-level acoustic features that enhance word recognition performance in the presence of noise. In contrast, background noise persists in the spiking activity of the high-resolution network, which results in a greater performance reduction across network layers.

**Figure 5.**
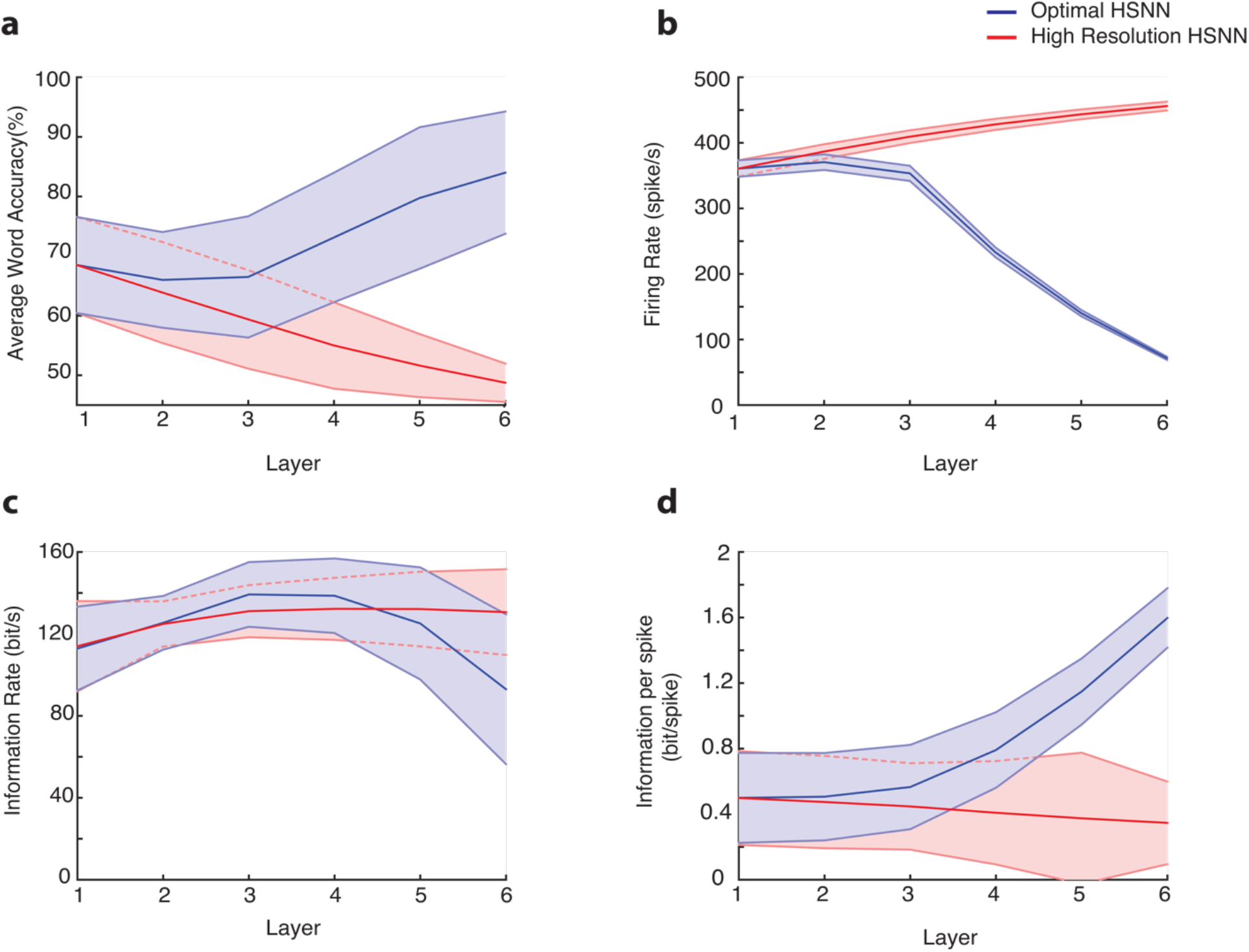
Hierarchical transformation between consecutive network layers enhances word recognition performance and robustness of the optimal HSNN. **(a)** The average word accuracy at 5 dB SNR systematically increases across network layers for the optimal HSNN (**a**, blue) whereas the high-resolution HSNN exhibits a systematic reduction in word recognition accuracy (**a**, red). For the high-resolution HSNN average firing rates (**b**, red), information rates (**c**, red), and information per spike (**d**, red) are relatively constant across layers indicating minimal transformations of the incoming acoustic information. In contrast, average firing rates (**b**, blue) and information rates (**c**, blue) both decrease between the first and last network layers of the optimal network, consistent with a sequential sparsification of the response and a reduction in the acoustic information encoded in the output spike trains. However, the information conveyed by single action potentials (**d**, blue) in the optimal HSNN sequentially increase between the first and last layer so that individual action potentials become progressively more informative across layers. Continuous curves show the mean whereas error contours designate the SD.

Although the classifier performance takes advantage of the hierarchical organization in the optimal HSNN, a similar trend is not observed for the transfer of acoustic information. First, firing rates decrease systematically across layers for the optimal HSNN, consistent with a sparser output representation (Fig. 5**b**, blue) as proposed for deep layers of the auditory pathway^2,20,33^. By comparison, firing rates are relatively stable across layers for the high-resolution network (Fig. 5**b**, red). We next measure the average mutual information (see Methods) in the presence of noise (5 dB) to identify how incoming acoustic information is sequentially transformed from layer-to-layer. For the optimal HSNN the information rates (i.e., bits / sec) decreases between the first and last layer (Fig. 5**c**, blue) whereas for the high-resolution network information is conserved across network layers (Fig. 5**c**, red). Thus, the layer-to-layer increase in word recognition accuracy observed for the optimal HSNN is accompanied by a loss of total acoustic information in the deep network layers. We next measure the average information conveyed by individual action potentials as way of determining how acoustic features are represented by individual precisely timed spikes. Surprisingly, the information conveyed by single action potentials is higher and increases across layers (Fig. 5**d**, blue). This contrast the high-resolution HSNN where information per spike remains relatively constant across layers (Fig. 5**d**, red). This indicates that individual action potentials become increasingly more informative from layer-to-layer in the optimal HSNN despite a reduction in firing rates. Taken together with the changes in spectro-temporal selectivity (Fig. 3), the findings are consistent with the hypothesis that the optimal HSNN produces a noise resilient sparse code in which invariant acoustic features are represented with isolated spikes. By comparison, the high-resolution network produces a dense response pattern that has a tendency to preserve incoming acoustic information, including the background noise and nonessential acoustic features, thus suffering in recognition performance.

We next asked whether the sequential layer-to-layer transformations of the optimal HSNN are required for robust coding of speech. Hypothetically, its plausible that similar performance could be achieved with a single layer network as long as each neuron accounts for the overall network receptive field transformations. To test this, we developed single-layer networks consisting of generalized linear model neurons^34^ with either a linear receptive field and Poisson spike train generator (LP network) or a linear receptive field and nonlinear stage followed by Poisson spike train generator (LNP network) (Fig. 6**a**; see Methods). The performance of the LP network, which accounts for the linear transformations of the optimal HSNN, was on average 21.7% lower than the optimal HSNN indicating that nonlinearities are critical to achieve high word recognition accuracy (Fig. 6**b**). Its plausible that this performance disparity can be overcome by incorporating a nonlinearity that models the rectifying effects in the spike generation process of neurons (LNP network). Doing so improves the performance to within 2.1% of the optimal HSNN when there is little background noise (SNR=20 dB, 85.6 % for optimal HSNN versus 82.5 % for LNP network). However, the performance degraded when background noise was added when compared to the optimal HSNN, with an overall performance reduction of 13.8 % at -5 dB SNR (58.4 % for optimal HSNN versus 44.6 % for LNP network).

**Figure 6.**
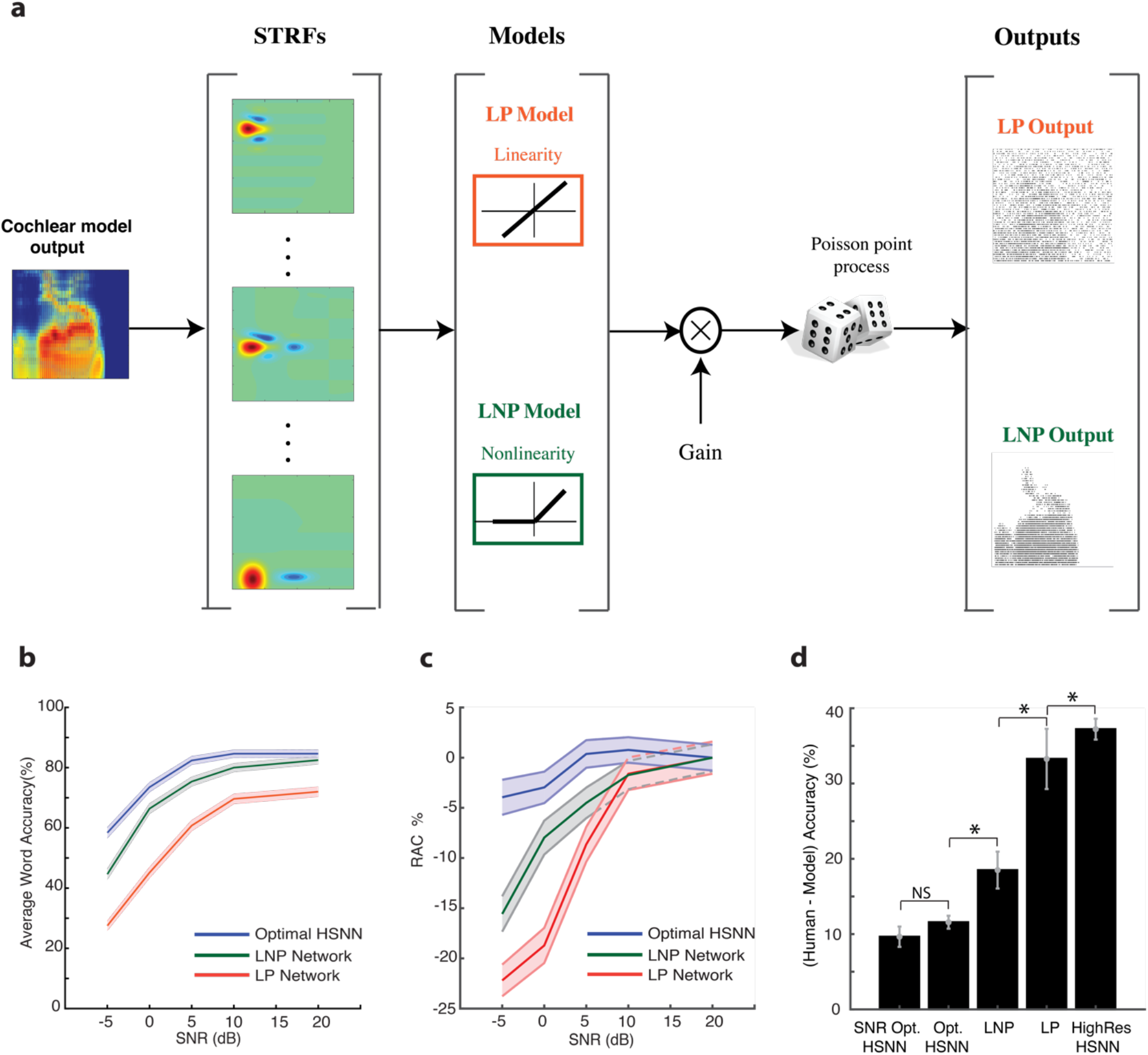
Optimal HSNN enhances robustness and outperforms single-layer generalized linear model networks with matched linear and nonlinear receptive field transformation. (a) Linear STRFs obtained at the output of the HSNN are used as to model the linear receptive field transformation of each neuron (see Methods). The LP network consists of an array of linear STRFs followed by a Poisson spike generator. The LNP network additionally incorporates a rectifying output stage following each STRF. (b) The optimal HSNN outperformance the LP network with an average performance improvement of 21.7% across SNRs. Nonlinear output rectification in the LNP network improves the performance to within 2% of the HSNN at 20 dB SNR. However, the average LNP performance was 7% lower than the optimal HSNN and performance degraded systematically with increasing noise levels (13.75 % performance reduction at -5 dB SNR) demonstrating enhanced robustness of the optimal HSNN. (c) The relative accuracy change (RAC=(A_model_-A_human_) − (A^20dB^_model_-A^20dB^ h_uman_)) was used to measure the divergence of each model across SNR when compared against human accuracy rates ^32^. An RAC of 0 across SNRs indicates that the model performance follows a similar noise robust trend when compared to humans. For the optimal HSNN, RACs were near zero across SNRs. RACs diverged substantially relative to human accuracy rates with increasing SNR for the LP and LNP networks. (d) Average accuracy difference between human and model data (A_human_ -A_model_). Average performance of the SNR optimal (optimized for each SNR) and optimal HSNN (optimized across all SNRs) are within ~10 % of the human word accuracy rates. The LNP (18.5 %), LP (33.3%) and high-resolution HSNN (37.2%) performance are substantially lower relative to humans. Asterisks designate significant differences (p<0.05, t-test with Bonferroni correction) and error bars designate SEM.

The robustness of each network was next examined by comparing the performance of each model against human performance trends. For each condition, we measured the relative accuracy change (RAC) between the model and human performance (Methods, Fig. 6**c**). The RAC of the optimal HSNN was near zero with a small reduction in RAC of only 3.9% at -5 dB SNR. Thus, the optimal HSNN follows a similar trend as humans across background noise levels. By comparison, both the LP and LNP performance diverged from human performance with increasing background noise with an overall RAC reduction of 22.2 % and 15.6% at -5 dB SNR, respectively. Thus, in contrast to the optimal HSNN trends which mirrors human data, the LP and LNP network performance diverged from the human trend with increasing background noise.

The average performance of each network was also compared against human word recognition accuracy. The accuracy for the optimal and SNR optimal HSNNs are not significantly differences when compared against human accuracy rates with an average reduction of 9.7% and 11.5%, respectively (p>0.05, t-test). Furthermore, the optimal HSNN outperformed all other models tested. The LNP, LP, and high-resolution HSNN exhibited a rank order reduction in performance relative to human accuracy (18.5 %, 33.3%, 37.2% respectively; p<0.05, t-test with Bonferroni Correction).

Overall, the findings indicate that although the linear and nonlinear receptive field transformations both contribute to the overall network performance, the sequential layer-to-layer transformations carried out by the optimal HSNN are critical for maintaining a noise robust representation that mirrors human performance trends.

### Optimal spiking timing resolution

Finally, we identified the spike timing resolution required to maximize recognition accuracy as previously identified when “reading out” neural activity in auditory cortex ^7,35^. To do so, we synthetically manipulating the temporal resolution of the output spike trains while measuring the word recognition accuracy at multiple SNRs (see Methods). An optimal spike timing resolution is identified within the vicinity of 4-14 ms for the optimal network (Fig. 7**a** and **b**) which is comparable to spike timing precision required for sound recognition in auditory cortex ^7,35^. By comparison, the high-resolution network requires a high temporal resolution of ~2 ms to achieve maximum word accuracy (46.6% accuracy across all SNRs; Fig. 8**c**), which is ~ 31.8% lower on average than the optimal network (78.4 % accuracy for the optimal HSNN across all SNRs). Taken across all SNRs, the optimal temporal resolution that maximized word accuracy rates is 6.5 ms, which is comparable to the spike timing resolution reported for optimal speech and vocalizations recognition in auditory cortex ^7,35^.

**Figure 7.**
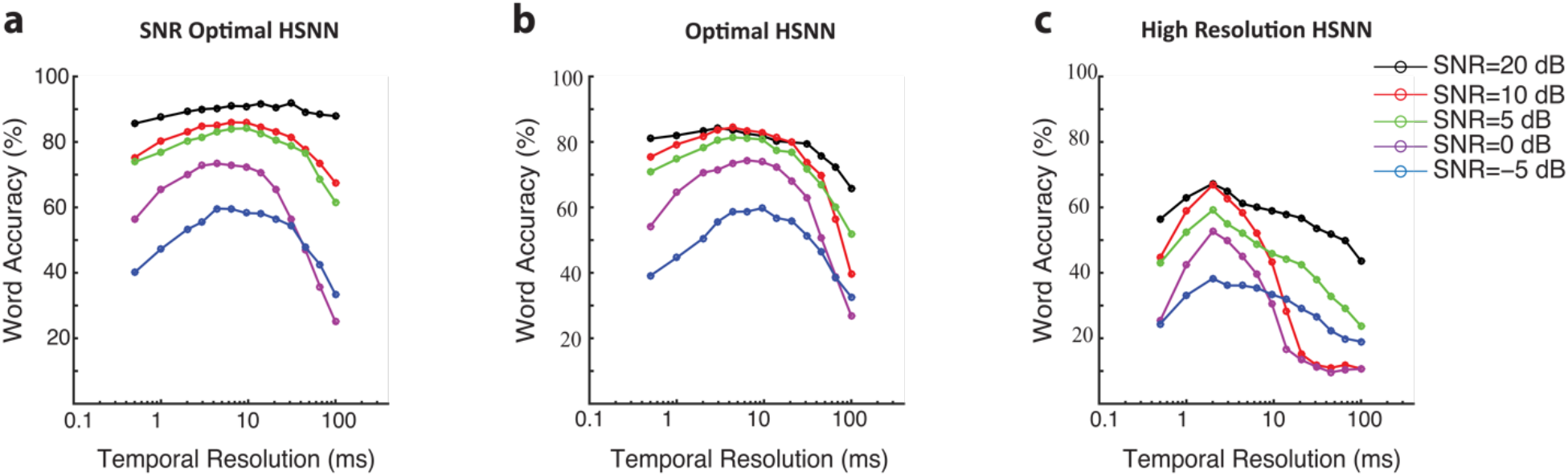
Optimal temporal resolution that maximize word recognition accuracy in noise. (**a**) Word accuracy rate as a function of spike train temporal resolution (bin widths 0.5-100 mms) and SNR (-5 to 20 dB) for the optimal (**a**) and high resolution networks (**c**). Each curve is computed by selecting the optimal scaling parameters for each SNR and measuring the word accuracy rate from the network outputs at multiple temporal resolutions. (**b**) Same as (**a**), except that global optimal scaling parameters were used for all SNRs tested. The temporal resolution that maximizes the word accuracy rate of the global optimal HSNN is 6.5 ms. (**c**) Word accuracy rate as a function of temporal resolution and SNR for the high-resolution network. The optimal temporal resolution for the high-resolution HSNN is 2 ms.

## Discussion

The results demonstrate that the hierarchical organization of the ascending auditory system is consistent with a near optimal strategy for feature extraction that maximizes sound recognition performance and is relatively impervious to noise. Upon optimizing the network organization on a behaviorally relevant word recognition task, the HSNN achieves high recognition accuracy and follows a similar noise robust trend that is within ~10% of human performance by sequentially refining the spectral and temporal selectivity from layer-to-layer. Similar noise robustness is not replicated with conventional receptive field based networks even when the receptive fields capture the linear integration of the optimal HSNN and a threshold nonlinearity was imposed. The sequential nonlinear transformations of the optimal HSNN preserve critical acoustic features for speech recognition while simultaneously discarding acoustic noise not relevant to the sound recognition task. These transformations mirror changes in selectivity along the ascending auditory pathway, including an extensive loss of temporal resolution^3^, slight loss of spectral resolution ^4-6^, and increase in sparsity ^2,20^. The simulations suggest that the orderly arrangement of receptive fields and sequential nonlinear transformations of the ascending auditory pathway may be critical to achieve a noise robust code.

Critical to our findings is the observation that the optimal network transformations described here are not expected a priori as a general sensory processing strategy and may in fact be unique to audition. For instance, changes in temporal selectivity between the retina, visual thalamus, and visual cortex are generally small and neurons in the visual pathway synchronize over a relatively narrow range of frequencies (typically < 20 Hz) ^36-39^. This differs dramatically from the observed increase in integration times reported here, systematic increase in synaptic potential time-constants^13,23-25^, and a corresponding reduction in synchronization ability^3^ observed between the auditory nerve and auditory cortex. By comparison, in the spatial domain, there is substantial divergence in connectivity between the retina and visual cortex since visual receptive fields sequentially grow in size between the periphery and cortex so as to occupy a larger area of retinotopic space ^40-42^. This contrasts changes in frequency receptive fields in which only a subtle increase in average bandwidth is observed between the auditory nerve and cortex^4-6,21,26^, consistent with findings from the optimal sound recognition strategy.

The findings outline a biologically plausible auditory coding strategy capable of efficiently achieving high recognition accuracy, particularly in the presence of noise. Although the auditory pathway is substantially more complex than the proposed HSSN, which lacks anatomical elements such as the binaural circuits in the brainstem and descending feedback, it is nonetheless surprising that the optimal strategy for speech recognition replicates sequential transformations observed along the auditory pathway. Furthermore, whereas auditory receptive fields can be more diverse than those of the HSNN, the receptive fields of the optimal HSSN nonetheless contain basic features seen across the auditory pathway including lateral inhibition, temporal inhibition or suppression, and sequentially increasing time-constants along the hierarchy ^6,26,43,45^. The HSSN employs several computational principles observed anatomically and physiologically, including the presence of spiking neurons, inhibitory connections, cotuning between excitation and inhibition, and a frequency specific localized circuitry, all of which likely contribute to its high performance. Furthermore, these sequential transformations appear to be critical since single layer generalized linear models designed to capture the overall transformations of the HSNN did not achieve comparable levels of performance.

Recent advances in deep neural networks (DNN) have made it possible to achieve high-levels of speech recognition performance approaching human performance limits^46,47^. Yet, these networks typically require tens-of-thousands of neurons and parameters to do so and the mechanisms leading to high recognition accuracy are based on neuron elements designed on principles of rate coding. The HSNN developed here, by comparison, employs temporal coding and organizational principles identified physiologically and approaches human performance levels with just 600 neurons and three meta-parameters that control the layer-to-layer transformations. Like the auditory pathway, the auditory HSNN is inherently temporal as it contains spiking neurons capable of precisely synchronizing to the sound features and exhibit hierarchical changes in time-scale across layers observed physiologically^3^. Furthermore, whereas DNNs rely on strictly excitatory connection weights between neuron, feature extraction in the HSNN is shaped by both excitatory and inhibitory circuitry as observed in central auditory structures ^10-13^. A challenge for future studies is to further reveal biologically realistic strategies for auditory signal processing, feature extraction, and classification, including descending feedback ^27^ and adaptive mechanisms ^1,29^, that together endow perceptual capabilities for sound recognition and promote robust coding.

## Materials and Methods

### Speech Corpus

Sounds in the experimental dataset consist of isolated digits (*zero* to *nine*) from eight male talkers from LDC TI46 corpus^22^. Ten utterances for each digit are used for a total of 800 sounds (8 talkers x 10 digits/subject × 10 utterances/digit). Words are temporally aligned based on the waveform onset (first upward crossing that exceeds 2 SD of the background noise level) and speech babble noise (generated by adding 7 randomly selected speech segments) is added at multiple signal-to-noise ratios (SNR=-5, 0, 5, 10, 15 and 20 dB). This range of SNR was selected to allow comparisons with human isolated word recognition performance in the presence of speech babble noise^32^.

### Auditory Model and Hierarchical spiking neural Network (HSNN)

We developed a multi-layer auditory network model consisting of a cochlear model stage containing gamma tone filters (0.1-4kHz; center frequencies 1/10^th^ octave separation; critical band resolution), envelope extraction and nonlinear compression^48^ followed by a HSNN as illustrated in Fig. 1. Several architectural and functional constraints are imposed on the spiking neural network to mirror auditory circuitry and physiology. First, the network contains six layers as there are six principal nuclei between the cochlea and cortex. Second, connections between consecutive layers contain both excitatory and inhibitory projections since long-range inhibitory projections between nuclei are pervasive in the ascending auditory system ^10,49^. Each layer in the network contains 53 excitatory and 53 inhibitory frequency organized neurons per layer which allows for 1/10^th^ octave resolution over the frequency range of the cochlear model (0.1-4 kHz). Furthermore, since ascending projections in the central auditory pathway are spatially localized and frequency specific ^18,49,50^, excitatory and inhibitory connection weights are modeled by co-tuned Gaussian profiles of unspecified connectivity width (Fig. 1**e**):

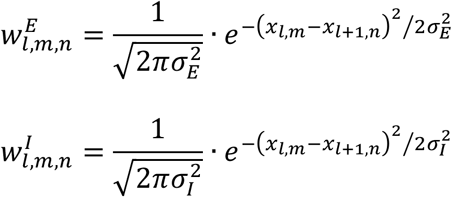

where 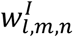 and 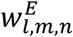 are the inhibitory and excitatory connection weights between the *m*-th and *n*-th neuron from layer *l* and *l+*1, *x_l,m_* and *x_l_*_+1_*_,m_* are the normalized spatial positions (0-1) along the frequency axis of the *m*-th and *n*-th neurons in layers *l* and *l*+1, and *σ_I_* and *σ_E_* are the inhibitory and excitatory connectivity widths (i.e., SD of Gaussian connection profiles), which determine the spatial spread and ultimately the frequency resolution of the ascending connections.

Each neuron in the network consists of a modified leaky integrate-and-fire (LIF) neuron ^51^ receiving excitatory and inhibitory presynaptic inputs (Fig. 1**e**). Given a presynaptic spike trains from the *m*-th neurons in network layer-*l* (*s_l,m_*(*t*)) the desired intracellular voltage of the *n*-th neuron in network layer *l*+1 is obtained as

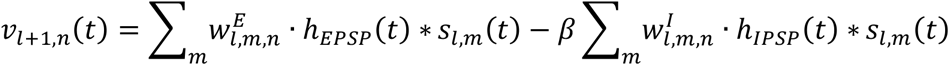

where * is the convolution operator, *β* is a weighting ratio between the injected excitatory and inhibitory currents, *h_EPSP_*(*t*) and *h_IPSP_*(*t*) are temporal kernels that model excitatory and inhibitory post synaptic potentials generated for each incoming spike as an alpha function (Fig. 1 **e**, red and blue curves)^51^. Since central auditory receptive fields often have extensive lateral inhibition/suppression beyond the central excitatory tuning area and inhibition is longer lasting and weaker ^5,6^ we require that *σ_I_* = 1.5 *· σ_e_*, *τ_I_ = 1.5 ·* τ_E_, and *β =* 2*/*3, as this produced realistic receptive field measurements. For simplicity, we use *σ* and *τ* interchangeably with *σ_E_* and *τ_I_*, since these determine the overall spectral and temporal resolution of each neuron.

Because the input to an LIF neuron is a current injection, we derived the injected current by deconvolving the LIF neuron time-constant from the desired membrane voltage

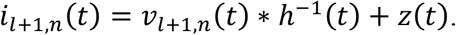

where *i_l_*_+1_*_,n_*(*t*) is the injected current for the *n*-th neuron in layer *l*+1 and *v_l_*_+1_*_,n_*(*t*) is the corresponding output voltage and *z*(*t*) is a noise current component. As we demonstrated previously ^19^, this procedure removes the influence of the cell membrane integration prior to injecting the current in the IF neuron compartment and allows us to precisely control the intracellular voltage delivered to each LIF neuron. Above 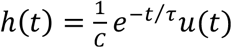 is the impulse response of the cell membrane (*u*(*t*) is the step function), C is the membrane capacitance, *τ*, is the membrane time-constant and *h^−^*^1^(*t*) is the inverse kernel (i.e., *h*(*t*) * *h*^−1^(*t*) *= δ*(*t*) where *δ*(*t*) is the Diract function). Because the EPSP time constant and the resulting temporal resolution of the intracellular voltage are largely influenced by the cell membrane integration, we require that *τ* = *τ_e_*. Finally, Gaussian white noise, *z*(*t*), is added to the injected current in order to generate spike timing variability (signal-to-noise ratio=15 dB) ^19^. Upon injecting the current, the resulting intracellular voltage follows *v_l_*_+1_*_,n_*(*t*) *+ z*(*t*) *\* h*(*t*) and the IF model generates spikes whenever the intracellular voltage exceeds a normalized threshold value^19^. The normalized threshold is specified for each network layer (*l*) as

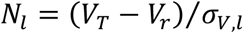

where *V_T_ =* −45 mV is the threshold voltage, *V_r_ =* −65 mV is the membrane resting potentials, and *σ_V,l_* is the standard deviation of the intracellular voltages for the population of neurons in layer *l*. As demonstrated previously, this normalized threshold represents the number of standard deviations the intracellular activity is away from the threshold activation and serves as a way of controlling the output sensitivity of each network layer. Upon generating a spike, the voltage is reset to the resting potential, a 1 ms refractory period is imposed, and the membrane temporal integration continues.

### Decision model

The neural outputs of the network consist of a spatio-temporal spiking pattern (e.g., Fig. 1**g** and **h**, bottom panels), which is expressed as a *N×M* matrix **R** with elements *r_n,i_* where *N*=53 is the number of frequency organized output neurons and *M* is the number of time bins. The number of time bins is dependent on the temporal resolution for each bin, ∆*t*, which is varied between 0.5 - 100 ms. Each response (*r_n,i_*; *n*–th neuron and *i* – th time bin) is assigned a 1 or 0 value indicating the presence or absence of spikes, respectively.

A modified Bernoulli Naïve Bayes classifier^52^ is used to read out the network spike trains and categorize individual speech words. The classified digit (*y*) is the one that maximizes posterior probability for a particular response according to

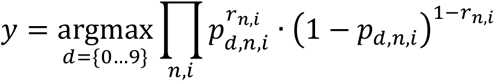

where *d=*0 … 9 are the digits to be identified, *p_d,n,i_* is the Bayesian likelihood, i.e. the probability that a particular digit, *d*, generates a spike (1) in a particular spatio-temporal bin (*n*-th neuron and *i*-th time bin).

### Network Constraints and Optimization

The primary objective is to determine the spectral and temporal resolution of the network connections as well as the network sensitivity necessary for robust speech recognition. Specifically, we hypothesize that the temporal and spectral resolution and sensitivity of each network layer need to be hierarchically organized across network layers in order to maximize speech recognition performance in the presence of noise. We thus optimize three key parameters, the time constant (*τ_l_*), connectivity widths (*σ_l_*), and normalized threshold (*N_l_*) that separately control these functional attributes of the network, where the index *l* designates the network layer (1-6). Given that spectro-temporal selectivity changes systematically and gradually between auditory nuclei, we constrained the parameters to vary smoothly from layer-to-layer according to the power law rules of Eqn. 1. The initial parameters for the first network layer, *τ*_1_ *=* 0.4 ms, *σ*_1_ *=* 0.0269 (equivalent to ~1/6 octave), and *N*_1_ *=* 0.5, are selected to allow for high-temporal and spectral resolution and high firing rates, analogous to physiological characteristics of auditory nerve fibers^3,4,26^ and inner hair cell ribbon synapse^23^. We optimize for the three scaling parameters *α, λ*, and *γ*, which determine the direction and magnitude of layer-to-layer changes and ultimately the network organization rules for temporal and spectral resolution and network sensitivity.

The optimization is carried using a cross-validation grid search procedure in which we maximized word accuracy rates (WAR). Initial tests are performed to determine a suitable search range for the scaling parameters and a final global search is performed over the resulting search space (*α =* 0.9 *−* 2.3*, λ =* 0.5 *−* 1.6 and *γ =* 0.8 *−* 1.5; 0.1 step size for all parameters). For each parameter combination, the network is required to identify the digits in the speech corpus with a ten-alternative forced choice task. For each iteration we select one utterance from the speech corpus (1 of 800) for validation and use the remaining utterances (799) to train the model by deriving the Bayesian likelihood functions (i.e., *p_d,n,i_*). The Bayesian classifier is then used to identify the validation utterances and compute WAR for that iteration (either 0 or 100% for each iteration). This procedure is iteratively repeated 800 times over all of the available utterances and the overall WAR is computed as the average over all iterations. This procedure is also repeated for five distinct signal-to-noise ratios (SNR=-5, 0, 5, 10, 20 dB). Example curves showing the WAR as a function of scaling parameters and SNR are shown in Fig. 2 (**a** and **b**, shown for 5 and 20dB). The global optimal solution for the scaling parameters is obtained by averaging WAR across all SNRs and selecting the scaling parameter combinations that maximize the WAR (Fig. 2**c**).

### Receptive Field and Mutual Information Calculation

To characterize the layer-to-layer transformations performed by the network, we compute spectro-temporal receptive fields (STRFs) and measure the mutual information conveyed by each neuron in the network. First, STRFs are obtained by delivering dynamic moving ripple sounds (DMR), which are statistically unbiased, and cross-correlating the output spike trains of each neuron with the DMR spectrotemporal envelope ^53^. For each STRF, we estimate the temporal and spectral resolution by computing the integration time and bandwidths, as described previously ^5^. Mutual information is calculated by delivering a sequence of digits (0 to 9) at 5 dB SNR to the network. The procedure is repeated 50 trials with different noise seeds and the spike trains from each neuron are converted into a dot-raster sampled at 2 ms temporal resolution. The mutual information is calculated for each neuron in the network using the procedure of Strong et al. ^54^ as described previously ^19^.

### Auditory System Data

Previously published data from single neurons in the auditory nerve (*n*=214) ^26^, auditory midbrain (Central Nucleus of the Inferior Colliculus, *n*=125)^48^, thalamus (Medial Geniculate Body, *n*=88) and primary auditory cortex (*n*=83)^6^ is used to quantify transformations in spectral and temporal selectivity between successive auditory nuclei. Using the measured spectro-temporal receptive fields of each neuron (Fig. 3), the spectral and temporal selectivity are quantified by computing integration times, response latencies, and bandwidths as described previously ^5^. Sequential changes in selectivity across ascending auditory nuclei are summarized by comparing the neural integration parameters of each auditory structure (Fig. 3**f-h**).

### Generalized Linear Model (GLM) Networks

To identify the role of linear and nonlinear receptive field transformations for noise robust coding, we developed two single-layers networks containing GLM neurons^34^ (Fig. 6**a**) that are designed to capture linear and nonlinear transformations of the HSNN.

First, we developed a single-layer LP (linear Poisson) network consisting of model neurons with linear spectro-temporal receptive fields followed by a Poisson spike train generator (Fig. 6**a**). For each output of the optimal network (*m*-th output) we measured the STRF and fitted it to a Gabor model (*STRF_m_*(*t, f_k_*))^43^. On average the fitted Gabor model accurately replicated the structure in the measured STRFs and on average accounted for 99% of the STRF variance (range 94-99.9%). The output firing rate of the *m*-th LP model neuron is obtained as

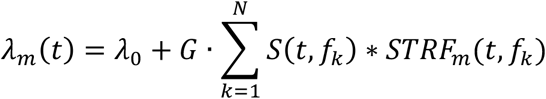

where *S*(*t, f_k_*) is the cochlear model output, * is the convolution operator, *G* is a gain term, and *λ*_0_ is required to assure that the spike rates are strictly positive and the firing maintains a linear relationship with the sound. *G* and *λ*_0_ are chosen so that the average firing rate taken across all output neurons and sounds matches the average firing rate of the optimal network and are strictly greater than zero. The firing rate functions for each channel, *λ_m_*(*t*), are then passed through a nonhomogenous Poisson point process in order to generate the spike trains for each output channel.

Next we explored the role of nonlinear rectification by incorporating a rectification stage in the LP model. The firing of the *m*-th neuron in the LNP (linear nonlinear Poisson) network is

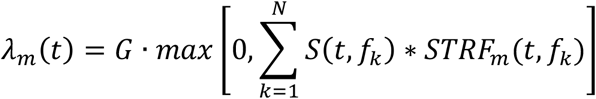

where the gain term, *G*, was chosen so that the average firing rate taken across all output neurons and all words matches the average firing rate of the optimal HSNN.

### Human Subject Data Comparison

Data was obtained from human subjects in an isolated monosyllabic word recognition task in the presence of speech babble noise ^32^. To enable comparison with the HSNN model conditions that we optimized for and tested (-5, 0, 5, 10, 20 dB SNR), human data (-6, -3, 0, 3, 6 dB SNR and quite) was fit to sigmoidal function and word accuracy rate values were estimated for human subjects at the model conditions tested. The sigmoid function fit accurate accounted for the human performance data with an average error of 0.9%. The average performance and trends with SNR of each model was compared against human performance set as a reference benchmark. The robustness of each model was also assessed by comparing how the word accuracy versus SNR trends deviate from human performance. The relative accuracy change RAC=(A_model_-A_human_) − (A^20dB^_model_-A^20dB^ _human_) was used to measure the divergence of each model across SNR when compared against human accuracy rates (i.e., Fig. 6**c**). An RAC of 0 indicates that the model performance follows a similar noise robust trend when compared to humans. Values <0 indicate that the model accuracy deviated (in units of %) from the human trend.

## Acknowledgments

We thank Heather Read and Ian Stevenson for providing feedback on the manuscript and E.D. Young for providing auditory nerve data. Research reported in this publication was partly supported by the National Institute On Deafness And Other Communication Disorders of the National Institutes of Health under Award Number R01DC015138 and a grant from the University of Connecticut Research Foundation. The content is solely the responsibility of the authors and does not necessarily represent the official views of the National Institutes of Health.

---

**Figure 1S.**
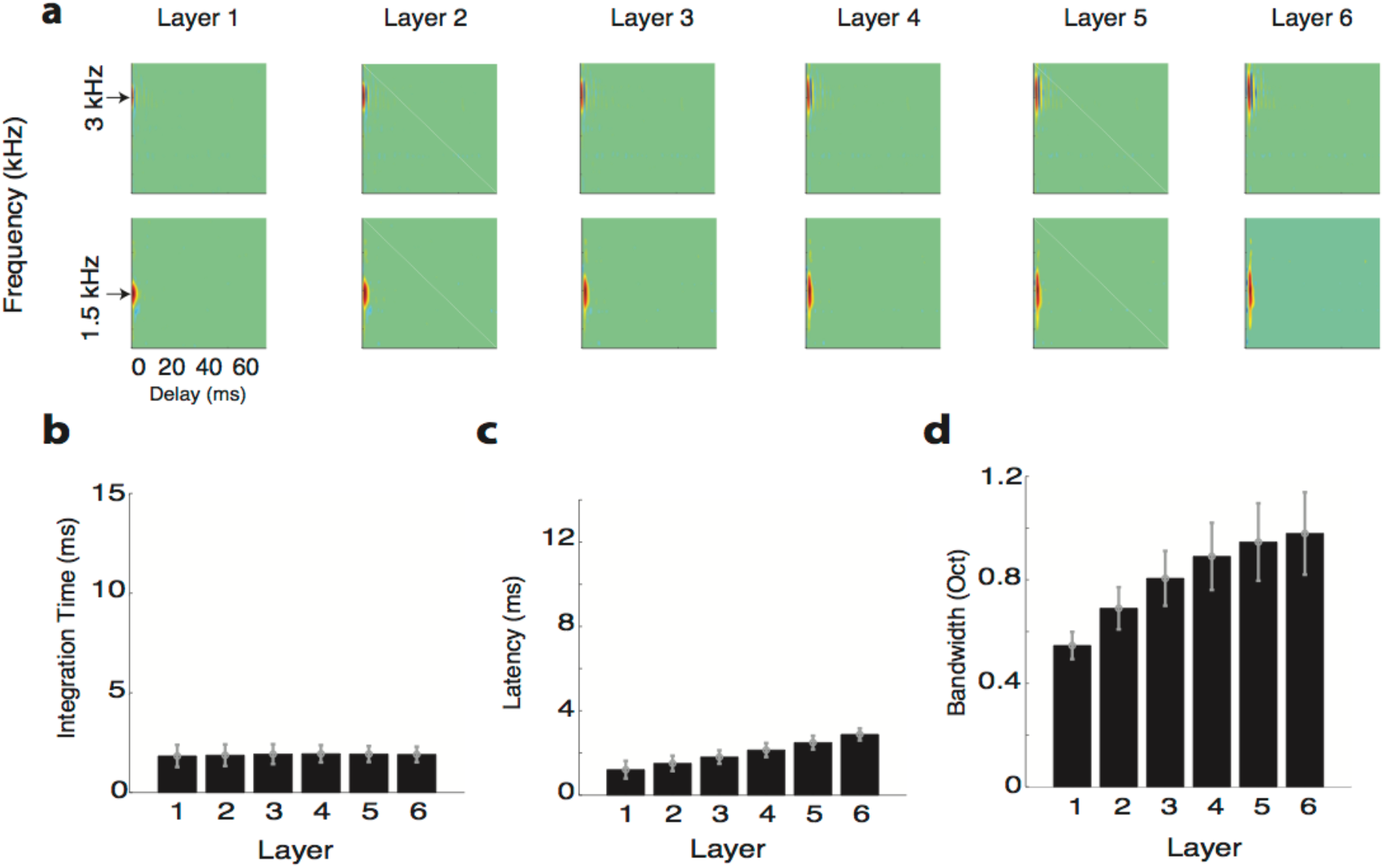
Receptive field transformations of the high-resolution network indicate that spectro-temporal information propagates with minimal processing across network layers. (**a**) Example spectro-temporal receptive field (STRF) measured for the optimal network maintain high-resolution and change minimally across network layers. Unlike the optimal network, the measured (**b**) integration times and (**c**) latencies change minimally and are relatively constant across the six network layers. (**d**) Bandwidths, by comparison, increase slightly across the six network layers and follow a similar trend as the optimal HSNN. The figure format follows the same convention as in Figure 3.

